# Predicting 30-day Hospital Readmissions Using Artificial Neural Networks with Medical Code Embedding

**DOI:** 10.1101/741504

**Authors:** Wenshuo Liu, Karandeep Singh, Andrew M. Ryan, Devraj Sukul, Elham Mahmoudi, Akbar K. Waljee, Cooper M. Stansbury, Ji Zhu, Brahmajee K. Nallamothu

## Abstract

Reducing unplanned readmissions is a major focus of current hospital quality efforts. In order to avoid unfair penalization, administrators and policymakers use prediction models to adjust for the performance of hospitals from healthcare claims data. Regression-based models are a commonly utilized method for such risk-standardization across hospitals; however, these models often suffer in accuracy. In this study we, compare four prediction models for unplanned patient readmission for patients hospitalized with acute myocardial infarction (AMI), congestive health failure (HF), and pneumonia (PNA) within the Nationwide Readmissions Database in 2014. We evaluated hierarchical logistic regression and compared its performance with gradient boosting and two models that utilize artificial neural network. We show that unsupervised Global Vector for Word Representations embedding representations of administrative claims data combined with artificial neural network classification models significantly improves prediction of 30-day readmission. Our best models increased the AUC for prediction of 30-day readmissions from 0.68 to 0.72 for AMI, 0.60 to 0.64 for HF, and 0.63 to 0.68 for PNA compared to hierarchical logistic regression. Furthermore, risk-standardized hospital readmission rates calculated from our artificial neural network model that employed embeddings led to reclassification of approximately 10% of hospitals across categories of hospital performance. This finding suggests that prediction models that incorporate new methods classify hospitals differently than traditional regression-based approaches and that their role in assessing hospital performance warrants further investigation.

## INTRODUCTION

Approximately 15% of patients discharged after an acute hospitalization are readmitted within 30 days, leading to potentially worse clinical outcomes and billions of dollars in healthcare costs [1]. Given these concerns, multiple quality efforts have been instituted in recent years to reduce readmissions in the United States. For example, the Medicare Hospital Readmission Reduction Program (HRRP) was created as part of the Patient Protection and Affordable Care Act and financially penalizes U.S. hospitals with excess 30-day readmission rates among Medicare beneficiaries [2,3]. Similar programs are being launched for patients with commercial insurance with the goal of further incentivizing hospitals to reduce readmissions [4,5].

Not surprisingly, the development of these programs has led to an increased demand for statistical models that accurately predict readmissions using available healthcare claims data. As the likelihood of readmission is related to key input features of patients (e.g., age and comorbidities), differences in the distribution of patients across hospitals based on such features may lead to unfair penalization of hospitals that care for more at-risk individuals. Therefore, using prediction statistical models to adjust for patient risk across hospitals is a major priority for accountability programs [6]. However, the performance of prediction models for readmissions have been generally poor. For example, existing methods that rely on regression-based models report area under the curve (AUC) for the receiver operating characteristic in the range of 0.63 to 0.65, suggesting limited discrimination for prediction [7,8]. Recent use of more flexible prediction models that leverage machine learning algorithms, such as random forest and traditional artificial neural network (ANN) models, have attempted to address this limitation with minimal improvements [9–11].

The purpose of this study was to explore whether advances in ANN models could improve prediction of 30-day readmission using administrative claims data and how this potential improvement may impact calculation of risk-standardized hospital readmission rates. ANN models abstract input features from large-scale datasets to predict output probability by approximating a combination of non-linear functions over the input future-space [12, 13]. Modern deployment of ANN models, including deep learning models, have been used successfully in a range of applications that include image classification and natural language processing [14–17], as well as prediction from electronic heath records [18,19]. We apply this approach to a large United States administrative claims data source focusing on 3 common conditions that were targeted under the HRRP: acute myocardial infarction (AMI), heart failure (HF) and pneumonia (PNA).

## METHODS

We conducted this study following the Transparent Reporting of a Multivariable Prediction Model for Individual Prognosis or Diagnosis (TRIPOD) reporting guidelines (see checklist in the Supplementary Information). All statistical code for replicating these analyses are available on the following GitHub repository: https://github.com/wenshuoliu/DLproj/tree/master/NRD. Data used for these analyses are publicly available at: https://www.hcup-us.ahrq.gov/tech_assist/centdist.jsp.

### Study Cohort

We used the 2014 Nationwide Readmissions Database (NRD) developed by the Agency for Healthcare Research and Quality (AHRQ) Healthcare Cost and Utilization Project (HCUP), which includes data on nearly 15 million admissions from 2048 hospitals [20–22]. The NRD has the advantage of including all payers, including government and commercial insurers. We identified patients hospitalized for AMI, HF, and PNA. We created a separate cohort for each condition using strategies for identifying patients that were adopted from prior published work [8, 23]. The cohort of index admissions for each condition was based on principal *International Classification of Diseases*-9 (ICD-9) diagnosis codes at discharge (e.g. in the case of AMI we used 410.xx, except for 410.x2) while excluding the following cases: (1) records with zero length of stay for AMI patients (n=4,926) per standards for constructing that cohort (as patients with AMI are unlikely to be discharged the same day); (2) patients who died in the hospital (n=13,896 for AMI, n=14,014 for HF, n=18,648 for PNA); (3) patients who left the hospital against medical advice (n=2,667 for AMI, 5,753 for HF, n=5,057 for PNA); (4) patients with hospitalizations and no 30-day follow up (i.e. discharged in December, 2014 (n=23,998 for AMI, n=44,264 for HF, n=47,523 for PNA)); (5) patients transferred to another acute care hospital (n=8,400 for AMI, n=5,393 for HF, n=4,839 for PNA); (6) patients of age < 18 years old at the time of admission (n=12 for AMI, n=409 for HF, n=28,159 for PNA); and (8) patients discharged from hospitals with less than 10 admissions (n=1,956 for AMI, n=1,221 for HF, n=418 for PNA). In circumstances where the same patient was admitted several times during the study period, we selected only the first admission. Flow diagrams for the cohort selection are shown in eFigure 1.

**Figure 1.**
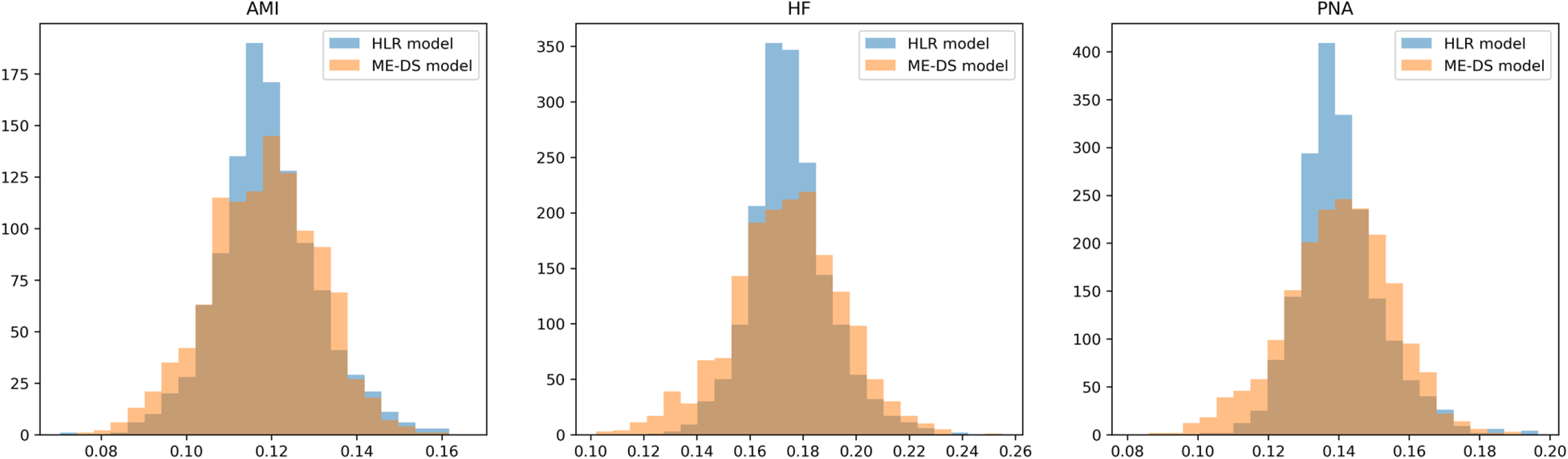
Distribution of risk-standardized hospital readmission rates for acute myocardial infarction (AMI), congestive health failure (HF), and pneumonia (PNA), generated by hierarchical logistic regression (HLR) model and the medical code embedding Deep Set architecture ANN (ME-DS) model.

### Study Variables

Our outcome was 30-day unplanned readmission created using the NRD Planned Readmission Algorithm [23]. The NRD also includes patient-level information on demographics and up to 30 ICD-9 diagnosis codes and 15 procedure codes from each hospitalization. Among the diagnosis codes, the principal diagnosis code at discharge represents the primary reason for the hospitalization while the rest represent comorbidities for the patient. To improve computational efficiency, we only included codes that appeared at least 10 times in the whole NRD database, reducing the number of ICD-9 diagnosis and ICD-9 procedure codes for inclusion in our analyses from 12,233 to 9,778 diagnosis codes and from 3,722 to 3,183 procedure codes, respectively.

### Statistical Models and Analysis

We evaluated four statistical models: 1) a hierarchical logistic regression model; 2) gradient boosting (using the eXtreme Gradient Boosting [XGBoost] [24] approach, a widely-used, decision tree-based machine learning algorithm) using ICD-9 diagnosis and procedure codes represented as dummy variables (1 if present, 0 if absent); 3) an ANN model using a feed-forward neural network with ICD-9 codes represented as dummy variables; and 4) an ANN model in which ICD-9 codes were represented as latent variables learned through a word embedding algorithm. We used hierarchical logistic regression as a baseline comparator given its ubiquitous use in health services and outcomes research. XGBoost is based on gradient boosted decision trees and it is designed for speed and performance. We used it given its rising popularity in recent years as a flexible machine learning algorithm for structured data. A more detailed explanation for the statistical models and ANN approaches as well as accompanying statistical code are available in the Supplementary Information.

The first model we constructed employed a hierarchical logistic regression model taking into account age, gender and co-morbidities. For co-morbidities, we used the well-established Elixauser Comorbitidy Index [25] to identify 29 variables to include as independent features in the model, with a hospital-specific intercept to account for patient clustering [7]. We implemented this model using the R package *lme4*.

For the second model, we applied XGBoost with the ICD-9 codes represented as dummy “1/0” input variables to provide a comparison from a popular machine learning tool. XGBoost has been well-recognized as an “off-the-shelf” algorithm that is highly effiecient and requires little hyper-parameter tuning to achieve state-of-the-art performance in a variety of tasks [26]. We implemented this model using the *Python* package *xgboost*.

For the third model, we implemented a feed-forward ANN model trained on dummy variable representation of ICD-9 diagnoses and procedure codes. The hospital ID was treated in the same way. We employed two hidden layers of neurons in this network to learn complex patterns from the input features instead of using human-engineered selection of variables (i.e., the Elixhauser Comorbidity Index). However, ANN models can be difficult to train because they require human parameter specification to reach optimal performance. Further, ANN models may be difficult to train due on large data-sets (over 4 million parameters in this case).

In the fourth model, we encoded 9,778 ICD-9 diagnosis and 3,183 procedure codes into 200- and 50-dimensional latent variable space, using the Global Vector for Word Representations (GloVe) algorithm [27]. We used GloVe, an unsupervised embedding algorithm to project ICD-9 co-occurrences to a lower dimension feature-space. The prescence of two ICD-9 diagnosis or procedure codes in a patient record during hospitalization was considered as a co-occurrence. We then counted the number of co-occurrences for each pair of ICD-9 diagnosis and/or procedure codes through the entire NRD database (excluding the testing set) and constructed the code embedding vectors according to the GloVe algorithm. A two-dimensional visualization of the embedding vectors of the ICD-9 diagnosis codes is shown in the eFigure 3. The visualization demonstrates that word embedding resulted in related diseases being closer to each other and is consistent with the application of word embedding algorithms in other administrative claims data [28, 29].

We used the deep set structure proposed by Zaheer et al [30] to incorporate ICD-9 diagnosis and procedure codes into the ANN model. This allowed us to account for varying counts of secondary ICD-9 diagnosis and procedure codes across patients and allow our model to be invariant to the ordering of these codes (e.g., the 2^nd^ and the 10^th^ code are interchangeable). The hospital ID was embedded into a 1-dimensional variable – conceptually this is similar to the hospital-level random intercept used in the hierarchical logistic regression models. The architectures of the two ANN models are shown in eFigure 2. The implementation of the ANN models was done using the *Python* packages *Keras* and *Tensorflow*.

To avoid the risk of overfitting, each of the study cohorts were divided into training, validation (for parameter tuning), and final testing sets at a proportion of 80%, 10%, and 10%, stratified by hospitals (i.e., within each hospital). We calculated AUC for the standard hierarchical logistic regression model, the XGBoost model and both ANN models on the final testing set, with the 95% confidence interval given from a 10-fold cross-validation. Once the models were developed, we then calculated risk-standardized hospital readmission rates for both the hierarchical logistic regression and ANN models (as the ANN models were superior to the gradient boosting model). We calculated these using predictive margin, which is a generalization of risk adjustment that can be applied for both linear and non-linear models (like ANN models) [31, 32]. Specifically, the predictive margin for a hospital is defined as the average predicted readmission rate if everyone in the cohort had been admitted to that hospital. Benefits of predictive margins over conditional approaches have been discussed in Chang et al [33]. We compared this approach to the traditional approach for calculating risk-standardized hospital readmission rates in hierarchical logistic regression models that uses the predicted over expected readmission ratio for each hospital and then multiplying by the overall unadjusted readmission rate [7]; importantly, we found similar results (see eFigure 4).

## RESULTS

### Study Cohort

Our study cohort included 202,038 admissions for AMI, 303,233 admissions for HF, and 327,833 admissions for PNA, with unadjusted 30-day readmission rates of 12.0%, 17.7% and 14.3% respectively. The mean (standard deviation) age was 66.8 (13.7) for AMI, 72.5 (14.2) for HF and 69.2 (16.8) for PNA, with the proportion of females 37.6%, 48.9% and 51.8%, respectively. Summary baseline characteristics are shown in Table 1 with additional details of the ICD-9 diagnosis and procedure codes in eTable 1. In these cohorts, we noticed an extremely skewed prevalence of ICD-9 diagnosis and procedure codes that were used to identify features for training related to comorbidities. For example, in the AMI cohort, three quarters of the 5,614 distinct secondary ICD-9 diagnosis codes appear less than 49 times (prevalence 0.02%), while the most frequent ICD-9 diagnosis code (i.e., 41.401 for coronary atherosclerosis of native coronary artery) appears 152,602 times (prevalence 75.5%). See eTable 1 for details.

**Table 1.** Summary statistics of the predictors for each of the 3 cohorts assessed in this study population.

|  | Acute Myocardial Infarction |  | Heart Failure |  | Pneumonia |  |
| --- | --- | --- | --- | --- | --- | --- |
|  | No Readmission | Readmission | No Readmission | Readmission | No Readmission | Readmission |
|  | N = 177,892 | N = 24,146 | N = 249,584 | N = 53,649 | N = 257,135 | N = 46,508 |
| Age, mean (std) | 66.3 (13.7) | 70.5 (13.3) | 72.5 (14.3) | 72.5 (13.9) | 68.6 (17.2) | 70.3 (15.8) |
| Female pct. | 36.60% | 45.00% | 48.80% | 49.30% | 52.60% | 50.20% |
| No. of diagnosis codes, mean (std) | 12.4 (6.1) | 15.7 (6.4) | 15.1 (5.5) | 16.2 (5.7) | 12.7 (5.8) | 14.7 (5.8) |
| No. of procedure codes, mean (std) | 5.6 (3.3) | 5.2 (3.9) | 1.1 (1.9) | 1.3 (2.1) | 0.7 (1.5) | 1.0 (1.8) |

### Performance of Prediction Models

Results of prediction of 30-day readmission as assessed by AUC are reported in Table 2 for each model and each cohort. The gradient boosting model utilizing XGBoost performed slightly better than the hierarchical logistic regression model and similar to the basic feed-forward ANN model. In general, the medical code embedding deep set architecture model generated the best results on all cohorts relative to the other three models. Compared with hierarchical logistic regression, the medical code embedding deep set architecture model improved the AUC from 0.68 (95% CI 0.678, 0.683) to 0.72 (95% CI 0.718, 0.722) for the AMI cohort, from 0.60 (95% CI 0.592, 0.597) to 0.64 (95% CI 0.635, 0.639) for the HF cohort, from 0.63 (95% CI 0.624, 0.632) to 0.68 (95% CI 0.678, 0.683) for the PNA cohort. In a sensitivity analysis, we repeated the same analysis on elderly patients (65 years old and above) and these are provided in eTable 2. Not unexpectedly, the overall AUCs decreased in the sensitivity analysis due to restriction of the cohort by age (which is a powerful predictor of readmission for patients); however, the margins for differences in AUCs across the four different statistical models increased slightly with this restriction by age.

**Table 2.** Results of prediction of 30-day readmission for acute myocardial infarction, heart failure and pneumonia as assessed by the Area under the curve of Receiver Operating Characteristic (AUC, with 95% confidence intervals in parentheses) given by the four models: the hierarchical logistic regression, XGBoost, feed-forward artificial neural network (ANN), and medical code embedding deep set architecture models.

| Methods | Acute Myocardial Infarction | Heart Failure | Pneumonia |
| --- | --- | --- | --- |
| Hierarchical Logistic Regression | 0.681 (0.678, 0.683) | 0.595 (0.592, 0.597) | 0.628 (0.624, 0.632) |
| XGBoost | 0.702 (0.698, 0.705) | 0.614 (0.611, 0.617) | 0.654 (0.651, 0.656) |
| Feed-Forward ANN | 0.707 (0.705, 0.709) | 0.623 (0.620, 0.626) | 0.663 (0.660, 0.666) |
| Medical Code Embedding Deep Set<br>Architecture | 0.720 (0.718, 0.722) | 0.637 (0.635, 0.639) | 0.680 (0.678, 0.683) |

### Risk-Standardized Hospital Readmission Rates

Given its overall higher performance, we compared risk-standardized hospital readmission rates calculated from the medical code embedding deep set architecture model with those calculated from the hierarchical logistic regression model. The histograms and summaries of these results are shown in Figure 1. Distributions of the risk-standardized hospital readmission rates from the two models were similar with just a modest shift downward in the mean for the medical code embedding deep set architecture model. We observed substantial differences in terms of rankings of individual hospitals between the two models. For both models, we divided the hospitals into three groups based on quintiles of predicted risk-standardized hospital readmission rates: top 20%, middle 60% and bottom 20%. For AMI, the medical code embedding deep set architecture model classified 72 (6.4%) hospitals in the middle 60% that the hierarchical model classified in the top 20% and classified 37 (3.3%) hospitals in the middle 60% that the hierarchical model classified in the bottom 20%. Results were similar for the HF and PNA cohorts (Table 3).

**Table 3.** Cross tabulation of divided groups between the hierarchical logistic regression (HLR) and the medical code embedding Deep Set architecture (ME-DS) model for each cohort.

|  | Acute Myocardial Infarction |  |  |  | Heart Failure |  |  |  | Pneumonia |  |  |  |
| --- | --- | --- | --- | --- | --- | --- | --- | --- | --- | --- | --- | --- |
|  | Rank in HLR model |  |  |  |  |  |  |  |  |  |  |  |
| Rank in ME-DS model | Top 20% | Middle 60% | Bottom 20% | All | Top 20% | Middle 60% | Bottom 20% | All | Top 20% | Middle 60% | Bottom 20% | All |
| Top 20% | 151 | 72 | 0 | 223 | 235 | 106 | 0 | 341 | 261 | 122 | 0 | 383 |
| Middle 60% | 72 | 563 | 37 | 672 | 106 | 854 | 66 | 1026 | 122 | 949 | 82 | 1153 |
| Bottom 20% | 0 | 37 | 186 | 223 | 0 | 66 | 275 | 341 | 0 | 82 | 301 | 383 |
| All | 223 | 672 | 223 | 1118 | 341 | 1026 | 341 | 1708 | 383 | 1153 | 383 | 1919 |

## DISCUSSION

In recent years, ANN models have shown advantages over traditional statistical models in a variety of medical tasks [18, 19]. Whether the application of such models to administrative claims data brings similar improvement in specific tasks related to prediction is worth exploring. This is especially important given the ubiquitous nature of claims data for assessing quality and hospital performance. In this paper, we applied ANN models towards the task of predicting 30- day readmission after AMI, HF, and PNA hospitalizations and compared it to existing approaches that use input features from classification systems that rely on expert knowledge like hierarchical logistic regression models as well as gradient boosting. Our findings suggest ANN models provide more accurate predictions of readmission and generate risk-standardized hospital readmission rates that vary from commonly used hierarchical logistic regression models.

There has been substantial work performed on constructing risk prediction models to predict readmissions after a hospitalization. The most frequent way these models are employed is through regression-based models that include age, gender and co-morbidities as input features [7]. For co-morbidities, ICD-9 diagnosis and procedure codes obtained from administrative claims data are used as input features to adjust for differences in individual patient risk in these models; however, not all of the thousands of potential ICD-9 diagnosis and procedure codes are included in the models and selecting which to incorporate is an important step. The selection has been based largely on expert input and empirical studies that have been used to generate fixed classification systems like the Hierarchical Condition Categories [34] or Elixhauser Comorbidity Index [25].

An advantage of ANN models is their ability as a statistical model to include thousands of features, as well as capture potential non-linear effects and interactions of these features. ANN models do not rely on human-generated classification systems but learn to automate extraction of relevant features from the data. Yet few studies to date have employed these models in administrative claims data. We believe a primary reason for this is that ANN models can be difficult to train due to the issues related to parameter optimization and memory consumption in the setting of a large number of parameters – sometimes in the order of millions. In the few studies that have used ANN models with administrative claims data [9, 35, 36], their use also may not have fully captured their full potential for risk prediction. For example, the use of binary “1/0” input features for ICD-9 diagnosis and procedure codes may ignore hidden relationships across comorbidities, limiting the ability of ANN models to improve on traditional hierarchical logistic regression or other methods like gradient boosting.

Of course, there has been some work on predicting readmissions using ANN models in the published literature. Futoma et al. implemented the basic architecture of feed-forward ANN models and showed modest advantages over conventional methods [9]. A number of researchers proposed to embed medical concepts (including but not limited to ICD-9 diagnosis and procedural codes) into a latent variable space to capture their co-relationships [28, 29, 37]; however, these investigators used this approach largely for cohort creation rather than predicting clinical outcomes or risk-adjustment. Krompass et al [36] used Hellinger-distance based principal components analysis [38] to embed ICD-10 codes and then built a logistic regression model using the embedded codes as input features. They found marginal improvements in prediction of readmissions over a feed-forward neural network but were restricted by their limited sample size. Choi et al. [35] designed a graph-based attention model to supplement embedding with medical ontologies for various prediction tasks, including readmission. However, their model did not explicitly consider the fact that the medical codes are permutation invariant. In this paper, we took advantage of a novel word embedding approach, Global Vector for Word Representations (GloVe) [27], as well as a new and recently proposed deep set architecture [30] to fully capture the inter-relationship and the permutation-invariant nature of the diagnosis and procedure codes. These choices – which were purposeful and driven by our intuition on the benefits of ANN models for this specific task – resulted in improved accuracy of prediction for readmission for a word embedding deep set architecture model across all 3 conditions.

Our study should be interpreted in context of the following limitations. First, although we found ANN models outperformed hierarchical logistic regression models, it is uncertain whether these improvements will justify their use more broadly as this requires consideration of other issues. For example, ANN models require large-scale data sources to train. Even though such data were available given the NRD for our current work, these are not always available. But the widespread availability and application of administrative claims data in assessing quality and hospital performance justifies the need to explore ANN models (and other approaches) further. Second, ANN models are computationally intensive and retain a “blackbox” feel with its findings difficult to understand and explain to users (similar to other models like gradient boosting). These issues may make it less attractive to policymakers and administrators when there may be a need to justify why performance is lacking in a public program (e.g., HRRP). Third, ANN models may not work for applications beyond 30-day readmission in these 3 common conditions. Work is needed to compare the performance of ANN models with traditional approaches for other outcomes (e.g., mortality), rare diseases, or populations (i.e., non-hospitalized patients).

In summary, ANN models with medical code embeddings have higher predictive accuracy for 30-day readmission when compared with hierarchical logistic regression models and gradient boosting, a widely used, decision tree-based machine learning algorithm. Furthermore, ANN models generate risk-standardized hospital readmission rates that lead to differing assessments of hospital performance when compared to these other approaches. The role of ANN models in clinical and health services research warrants further investigation.

## Author Contributions

Wenshuo Liu: Study design, methodology development, data curation and statistical analysis, drafting of manuscript, critical review of manuscript.

Karandeep Singh: critical review of manuscript. Andrew M. Ryan: critical review of manuscript.

Devraj Sukul: data curation, critical review of manuscript. Elham Mahmoudi: critical review of manuscript.

Akbar K. Waljee: critical review of manuscript. Cooper M. Stansbury: critical review of manuscript.

Ji Zhu: Study design, methodology development, statistical analysis, drafting of manuscript, critical review of manuscript.

Brahmajee K. Nallamothu: Study design, data interpretation, drafting of manuscript, critical review of manuscript.

## Author Access to Data and Data Analysis

Wenshuo Liu and all study co-authors had full access to all the data in the study and takes responsibility for the integrity of the data and the accuracy of the data analysis

## Transparency Declaration

Wenshuo Liu, Ji Zhu, and Brahmajee Nallamothu affirm that this manuscript is an honest, accurate, and transparent account of the study being reported; that no important aspects of the study have been omitted; and that any discrepancies from the study as planned have been explained.

